# Effect of MRI Defacing on EEG Forward and Inverse Modeling

**DOI:** 10.1101/2025.03.11.642561

**Authors:** Angela Mitrovska, Stefan Haufe

## Abstract

Recent years have seen improvements in facial recognition software, which has increased the risk of reidentification of patients’ structural neuroimaging scans, e.g. obtained from magnetic resonance imaging (MRI). Thus, an anonymization procedure known as defacing has become the norm when publicly sharing patients’ scans. Defacing removes some of the facial features from the data, making it improbably to re-identify a patient from a 3D rendering of the image. However, certain tasks, such as localization of the sources of electroencephalographic (EEG) signals, require the creation of individual electrical volume conductor models of the human head from structural MRI data. Defacing could affect the co-registeration of MRI and EEG sensor positions and, more importantly, the model of electrical current flow itself. This study quantifies and maps the effect of defacing on individual volume conductor models and the localization accuracy of inverse solutions based on these models in a sample of ten subjects with known ground-truth (non-defaced) anatomy. Boundary and finite element modeling approaches (B/FEM) are compared.

**Clinical relevance:** This study enables clinicians, e.g. epileptologists, and neuroscientists to gauge the EEG source localization errror that is incurred due to using defaced MRI data for forward model construction, and identifies the regions most prone to mislocalization. Based on the results of the study, it is recommended to construct EEG volume conductor models before defacing if accurate source localization is desired.

## I. INTRODUCTION

The growth of the medical research field has been supported by data sharing and making data publicly available. Publicly available data plays an important role in studying particular diseases and conditions as well as in aiding new discoveries. However, regulations such as the General Data Protection Regulation (GDPR) [1] and the Health Insurance Portability and Accountability Act (HIPAA) [2], make it harder for researchers to have access to diverse data. Sharing patient data publicly poses ethical, legal, and technical challenges. For data to be publicly shared, HIPAA requires that full-face photographic images must be removed from the data [3], among other provisions. With the evolution of 3D rendering software, it has become possible to reconstruct detailed images showing facial anatomy. Thus, in the medical field, anonymization has emerged as a method enabling the protection of patient’s privacy. Almost all data-sharing neuroimaging initiatives use defacing to remove facial features before sharing the data [4]. The two most common defacing techniques are: removing all facial features (e.g., using FreeSurfer [5]) and blurring the face [4], [6].

Defacing by removing all facial features typically encompasses removing the internal facial features, i.e., eyes, nose, and mouth, from the images, as these features are considered particularly relevant when trying to recognize an individual [7]. Having morphological analyses of the brain’s structure as the main purpose of the MRI acquisition in mind, care is typically taken to ensure that only non-brain tissue is removed, while brain tissue is kept intact [7]. In theory, the defacing process only affects voxels that are part of the face of the individual; however, their removal has been shown to also affect the automatic quantification of gray and white matter structures [8]. This could be due to the fact that segmentation algorithms use the face for context.

A less often considered side effect of defacing is that it could be detrimental in tasks that rely on facial areas as anatomical landmarks when working with other types of neuroimaging data. One of these tasks is source localization of magneto- and encephalography (M/EEG) signals, where the face is a useful anchor point when co-registering structural MRI with the location of EEG or MEG sensors [3]. The accuracy of the co-registration is vital when performing source localization and could be affected by the process of defacing. More importantly, neural return currents and magnetic fields sensed by EEG and MEG traverse through the facial region. Omitting that region is thus expected to affect volume conductor models and the localization of sources based thereon. This effect is hypothesized to occur predominantly in the frontal brain regions.

The paper is organized as follows. Section II presents an overview of forward and reverse M/EEG modeling and introduces the different types of forward models. Section III provides an overview of the method used in the study. In Section IV, the obtained results are presented and discussed. Finally, Section V concludes the paper.

## II. FORWARD AND INVERSE MODELING

Brain imaging technologies such as EEG allow researchers and clinicians to track neuronal populations with millisecond precision. However, to establish a relation between the neuronal currents generated in the structures of the brain and the voltage potentials measured at electrodes located on the scalp, an electrical volume conductor model of the human head is required [9]. This results in a so-called lead field matrix or forward model, which is the basis for EEG inverse modeling.

Volume conductor models can be obtained using the boundary element method (BEM) or the finite element method (FEM) [9]. As BEM modeling is less computationally demanding, it has become the predominant approach in EEG source imaging. In BEM modeling, only the major tissues such as the brain, skull, and scalp are represented by tissue boundaries derived from the individual’s anatomy [9]. However, the cerebrospinal fluid (CSF) is often not included, and the boundaries must entirely enclose each other, limiting the realism and accuracy [9]. On the other hand, FEM allows for the encoding of more anatomical details. FEM models most often include the gyri/sulci of the cortex, the thin layer of CSF, and the small but delicate structures of the skull [9], [10]. However, FEM can be potentially computationally costly [10].

This study investigates both BEM and FEM modeling to assess how the finer anatomical details in FEM models are impacted by the defacing process.

## III. METHODS

The MRI data for the study were taken from the Information eXtraction from Images (IXI) dataset [11]. IXI contains nearly 600 MRIs from normal, healthy, subjects. Ten subjects were randomly selected. All chosen scans are T1-weighted MRI images acquired at 3T field strength. Importantly, the scans were provided without any facial anonymization and were neither co-registered nor normalized to a common space.

The FreeSurfer `recon-all` pipeline [12] was applied to the raw image data to extract the cortical surface. Next, a defaced dataset was generated in addition to the original data. To achieve this, all voxels from the subjects’ facial areas were removed using FreeSurfer’s `mri-deface` function.

All further analyses were conducted using Matlab (The MathWorks, Inc.) separately for defaced and non-defaced data. Data were loaded into Brainstorm [13], where three BEM surfaces representing the inner skull, outer skull, and outer head (skin) surfaces were extracted from the MR image (defaced or not) and pre-extracted cortical surface data. Tetrahedral FEM meshes for the same three compartments were generated using iso2mesh [14]. An example illustrating the effect of facial feature removal on the outer head surface is given in Figure 1.

**Fig. 1.**
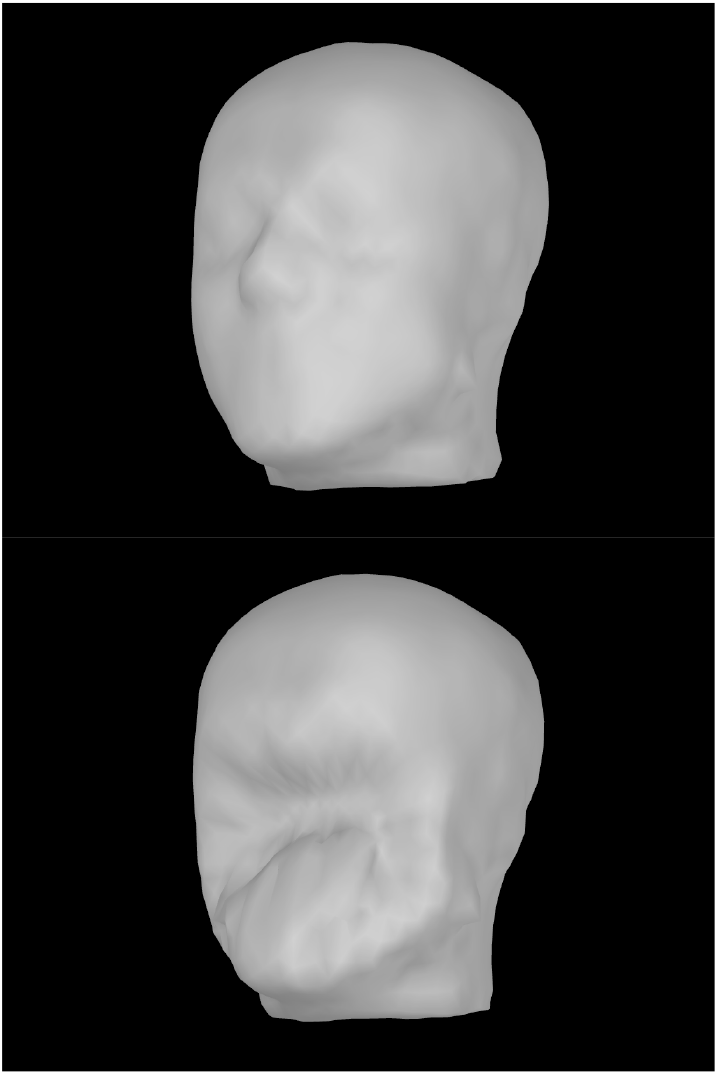
Non-defaced compared to defaced head anatomy.

A total of 65 EEG sensors were modeled at default positions according to the international 10-10 placement system [15]. Both BEM and FEM were used to compute the lead fields for 15002 source locations covering the entire cortical surface for each head model. BEM modeling was conducted using the OpenMEEG package [16], while FEM modeling was carried out using the DUNEuro package [17]. Default tissue conductivities (brain, skull, skin) provided by Brainstorm were used. Since the defacing was carried out after the FreeSurfer preprocessing, no alignment was required between the source space of the defaced and non-defaced anatomies for each individual; both remained identical. Thus, the comparison of results obtained on defaced and original data relied on identical brain voxels and sensor locations.

Quantitive comparisons of head models were performed following [9]. For each individual, the lead fields computed from the non-defaced anatomy served as the ground truth. The lead fields from the defaced anatomy were then correspondingly compared to the ground truth, to quantify the impact of defacing. The comparison was conducted using: (1) the strength of the predicted scalp potentials for identical sources, (2) the correlation of the lead fields for identical sources, computed using Matlab’s subspace command, and (3) the Euclidean distance between the estimated source locations in the defaced models and their corresponding ground-truth locations in the non-defaced head models.

The lead fields generated using the defaced anatomy were compared to those obtained using the non-defaced anatomy for each of the ten subjects. The *M ×* 3 lead fields of the non-defaced and defaced model at the *i*-th cortical location, *r*_*i*_ are referred to as 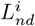 and 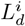, repectively, where *M* is the number of electrodes (in this case *M* = 65).

Following [9], the quantitative evaluation presented here is entirely based on measures that are invariant to rotations in 3D space. First, lead fields were compared based on the relative strength of their resulting scalp potentials. The relative lead field strength (gain) at a cortical location is defined as

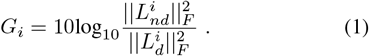

Since gain is expressed as the logarithm of a power ratio, it is measured on a *dB* scale. *G*_*i*_ is independent of the orientation of the source currents, as presented in [9].

Second, the correlation between the lead fields was computed based on the largest principle angle between the subspaces spanned by 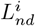 and 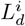. This was computed using Matlab’s subspace command, and, just as the gain, it is invariant to rotations in 3D space. The subspace angle is independent of any scaling, making it a suitable measure of subspace correlation [9]. Subspace angles were further normalized to the interval [0, 1], where 1 stands for orthogonal lead fields, and 0 stands for lead fields identical up to arbitrary linear transformation. Subspace correlation is defined as (1 - subspace angle) and is higher for more similar lead fields.

Finally, an EEG source reconstruction was simulated to assess the effect of defacing. Scalp potentials were generated using the ground truth model for all individuals, while localization was performed in the defaced model. The simulation was carried out separately for each cortical location. In the *i*- th run of the simulation, the lead field 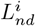 at the location, *r*_*i*_ was projected onto the normal vector of the cortical surface at *r*_*i*_, *n*_*i*_, leading to a single M-dimensional vector 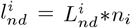. This vector represents the scalp potential that would be generated in the subject’s original anatomy by a dipolar current source at *r*_*i*_ oriented perpendicular to the cortical surface and was treated as a pseudo-EEG measurement. Localization was performed by sweeping through all cortical locations *h*_*j*_ in the defaced head model and comparing lead fields using the subspace correlation criterion. The location *r*_*j*_ with the highest subspace correlation, *h*^*max*^, was identified as the estimated source location in the defaced model. The Euclidean distance between *r*_*i*_ and the estimated source location *r*_*j*_, measured in millimeters, quantified the EEG localization accuracy, with larger distances indicating larger localization errors.

## IV. RESULTS AND DISCUSSION

Head models were quantitatively compared using three performance metrics (gain, subspace angle, and localization error) across two types of forward models (BEM and FEM). Each of the six resulting comparisons is presented in a separate plot. While the subspace angle is a symmetric measure, gain and localization error calculations assume that the individual non-defaced model is the ground truth, and the comparisons were carried out with respect to it.

From Figure 2, it can be observed that, for the BEM models, the range of gain is relatively narrow, and for most of the subjects it extends from 0 to 1 dB. However, for subject 9, the range of gain extends from -6 to -3 dB. For most of the subjects, there is an underestimation of the lead field in the frontal lobe when using defaced data to compute the head model. In contrast, subject 9, exhibits an overestimation of the lead field in all cortical areas, while subject 1 shows an overestimation in the frontal, parietal and occipital lobes.

**Fig. 2.**
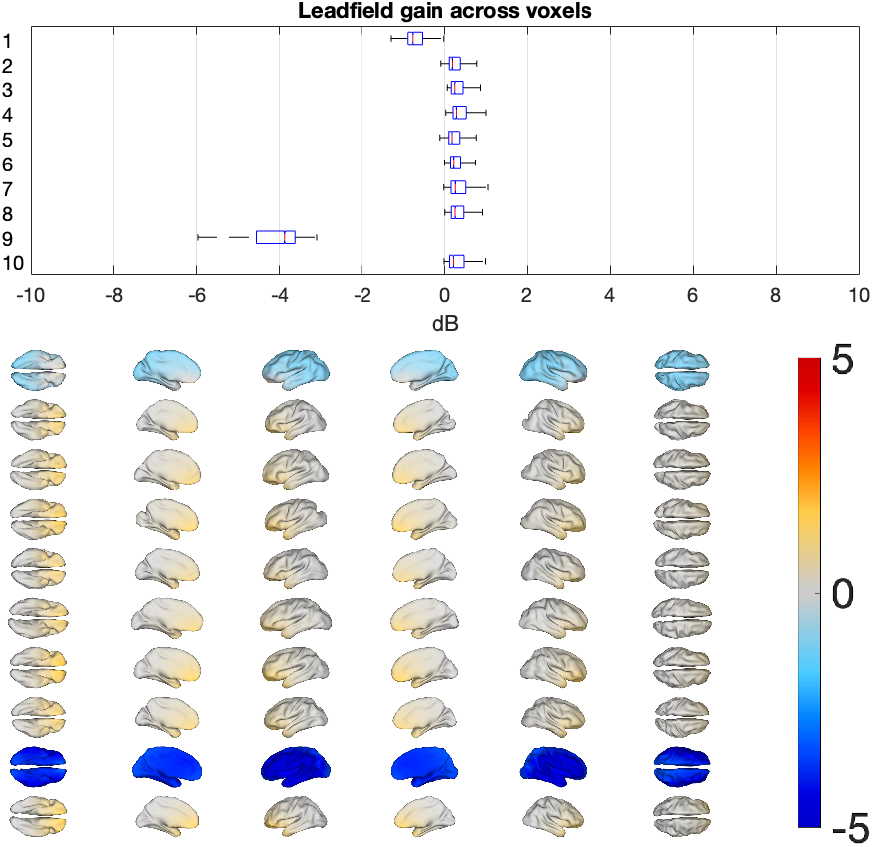
Lead field gain observed across all cortical locations when comparing defaced to non-defaced BEM models. Values closer to zero indicate better approximation performance. Upper panels: Median, 25th and 75th percentile, and most extreme values attained across all cortical locations. Lower panels: Six views of the topographical distributions of the gain for all ten subjects.

The results for the FEM models can be observed in Figure 3. For the FEM models, the gain range is wider, and the distribution of values is far less uniform across subjects. However, the results are mostly consistent with the observed values for the BEM models. For subject 9, there is an overestimation of the lead field strength across all of the lobes. For most other subjects, there is an underestimation of the lead field strength in the superficial frontal and central lobes.

**Fig. 3.**
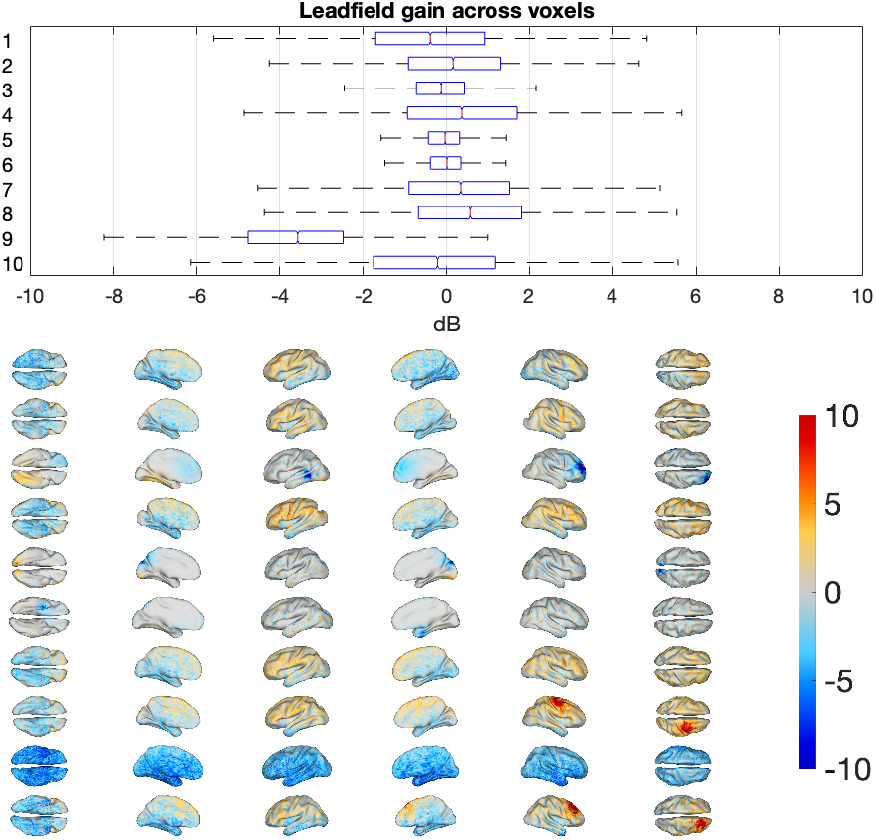
Lead field gain observed across all cortical locations when comparing defaced to non-defaced FEM models. Values closer to zero indicate better approximation performance. All graphs analogous to Figure 2; see caption for detail.

Figure 4 depicts the subspace correlation results for the BEM models. It can be observed that the correlation is quite high, except for subject 9, where the correlation overall is the lowest. The worst correlation is achieved in the frontal lobe, which is likely the most affected by the process of defacing.

**Fig. 4.**
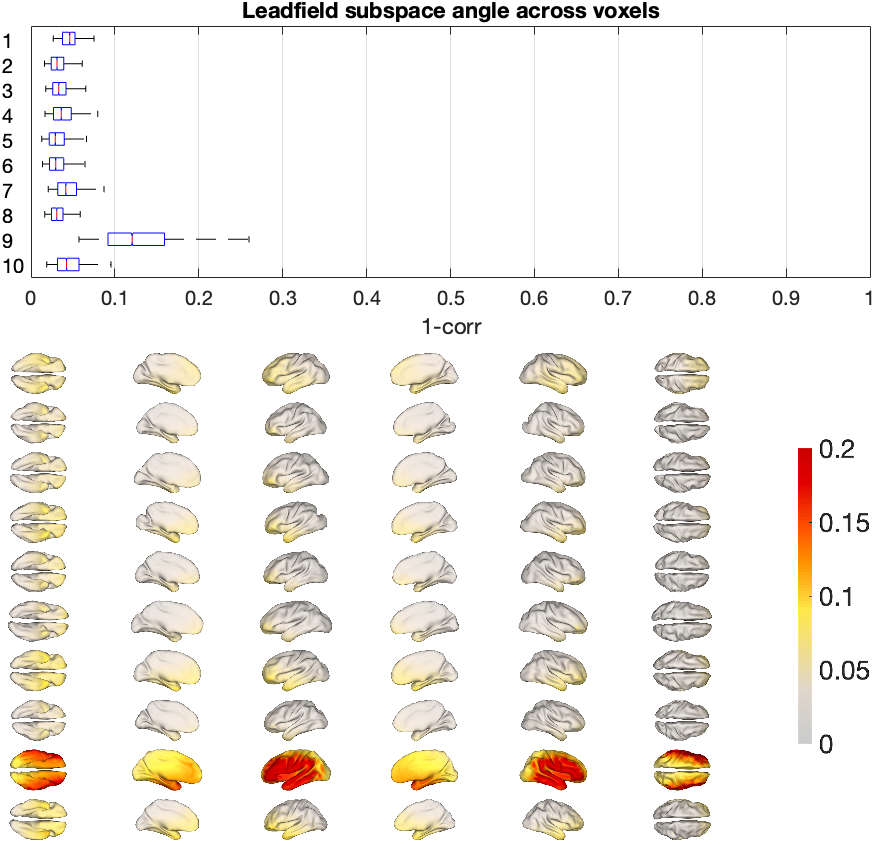
Subspace angle (1-subspace correlation) achieved across all cortical locations when comparing defaced to non-defaced BEM models. Smaller values indicate better approximation performance. All graphs analogous to Figure 2; see caption for detail.

From Figure 5, it can be observed that for the FEM models, the correlation is much lower than for the BEM models. This can be attributed to the fact that FEM models are more detailed than BEM models. For the FEM models, the worst correlation is once again achieved in the frontal and central lobes.

**Fig. 5.**
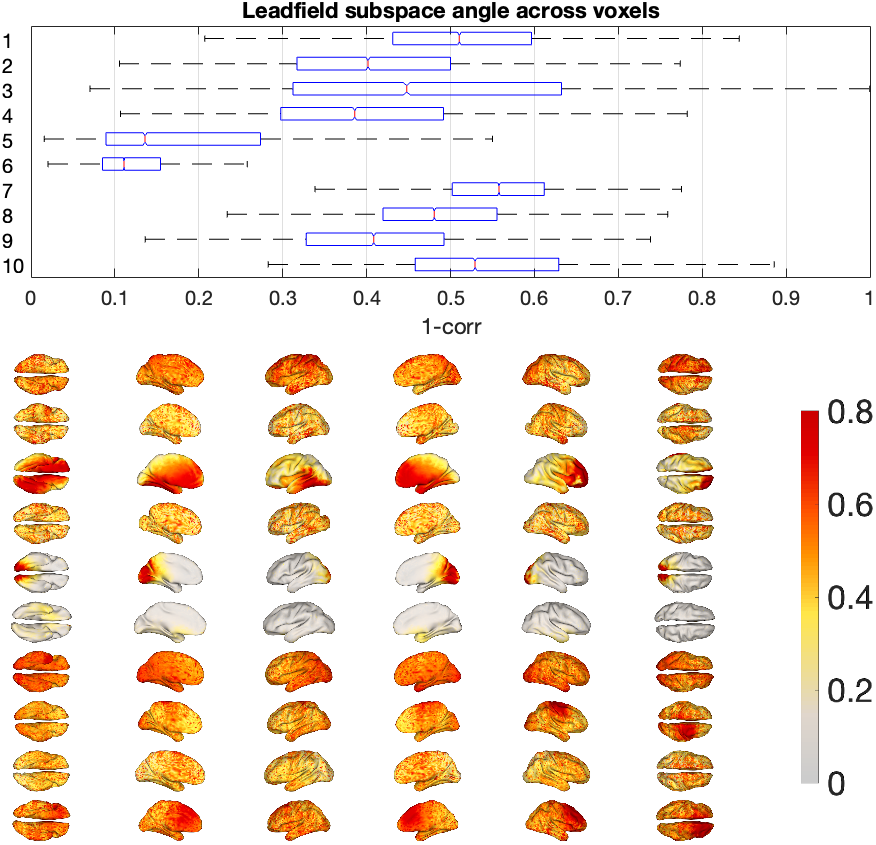
Subspace angle (1-subspace correlation) achieved across all cortical locations when comparing defaced to non-defaced FEM models. Smaller values indicate better approximation performance. All graphs analogous to Figure 2; see caption for detail.

In Figure 6, the results for the simulated EEG source localization in the BEM models are presented. The errors incurred for all subjects range between 0 and 12 mm. The highest localization error is achieved in subject 6.

**Fig. 6.**
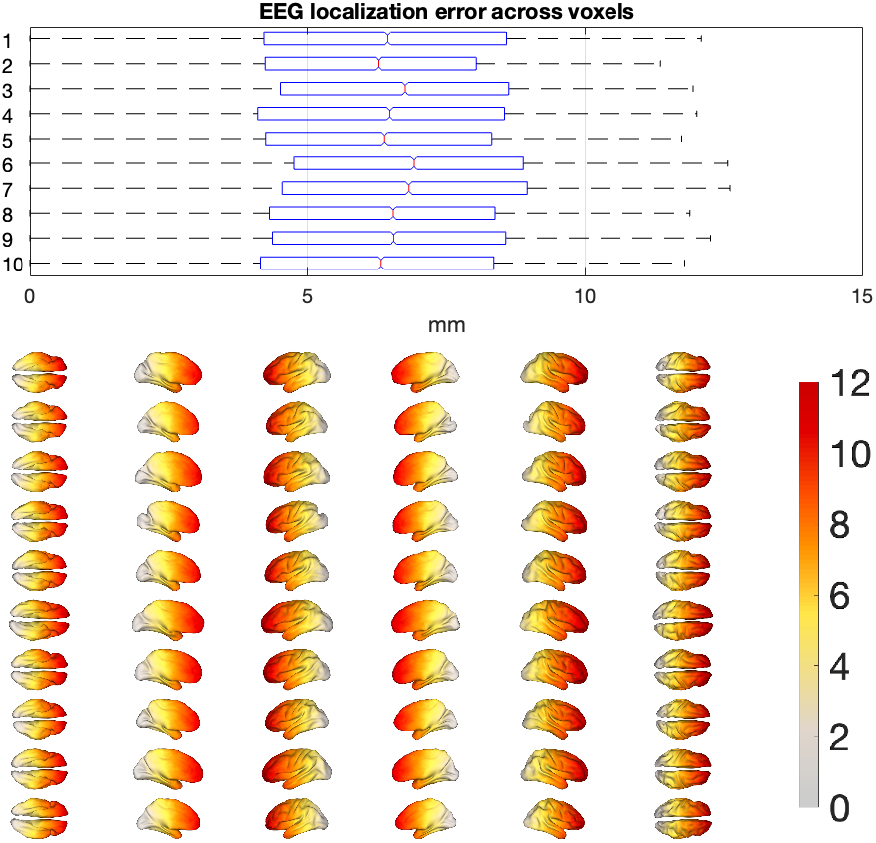
Localization error incurred using BEM models for dipolar sources placed across all cortical locations when performing EEG source imaging in the defaced head model. All graphs analogous to Figure 2; see caption for detail.

Figure 7 presents EEG source localization results for FEM modeling. The results for all subjects range between 0 and 26 mm, where the highest error is once again achieved in subject 6. The topographical distribution of the localization errors does not necessarily resemble the lead field subspace correlations shown in the previous figures, which is a finding that should be explored further.

**Fig. 7.**
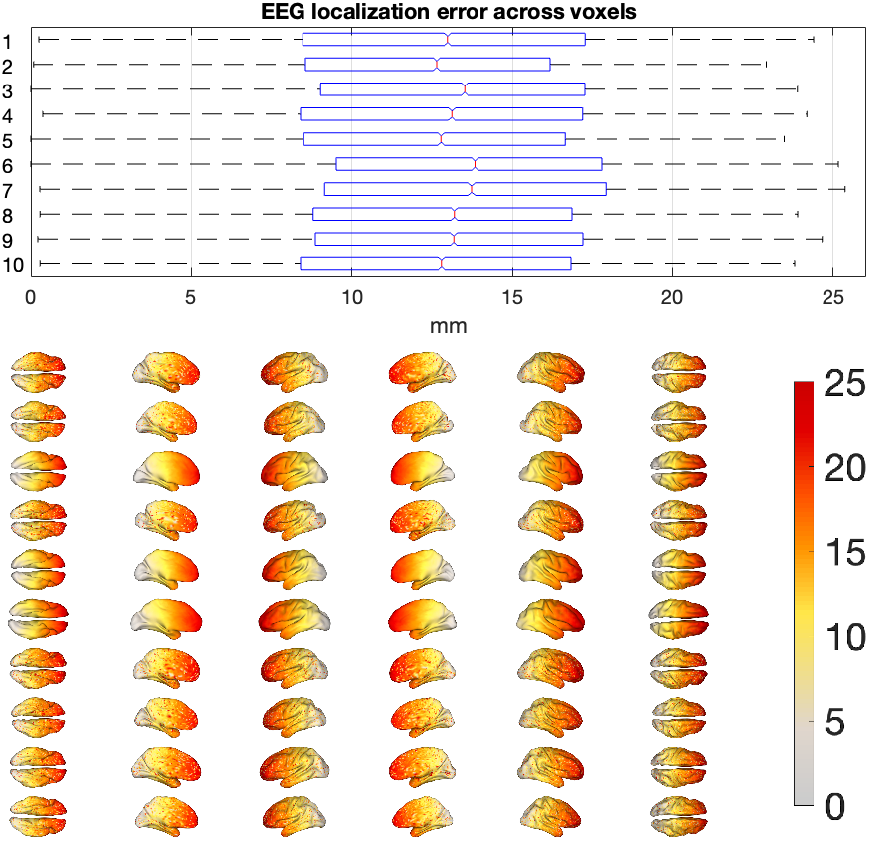
Localization error incurred using FEM models for dipolar sources placed across all cortical locations when performing EEG source imaging in the defaced head model. All graphs analogous to Figure 2; see caption for detail.

From the overall results, it can be concluded that the BEM models are not affected by the process of defacing to a great degree, with the exception of subject 9. A possible explanation is presented in Figure 8, which shows that defacing in subject 9 resulted in an unintended expansion of the external head anatomy, impacting cortical regions. In contrast, for most subjects, the defaced anatomy closely resembles the original, apart from the facial region, as presented in Figure 9. This study highlights the significance of edge cases like subject 9, where defacing-induced alterations could potentially distort results when analyzing specific conditions across individuals.

**Fig. 8.**
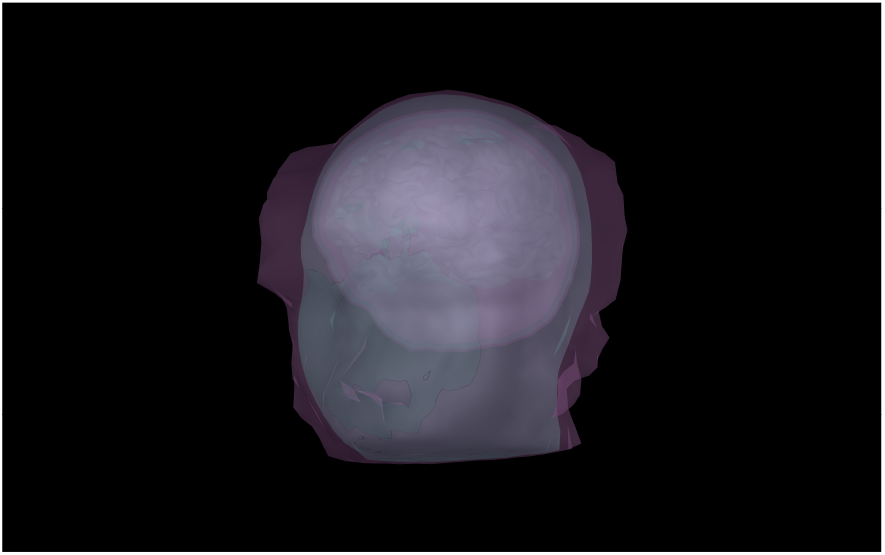
Defaced and non-defaced anatomy of subject 9.

**Fig. 9.**
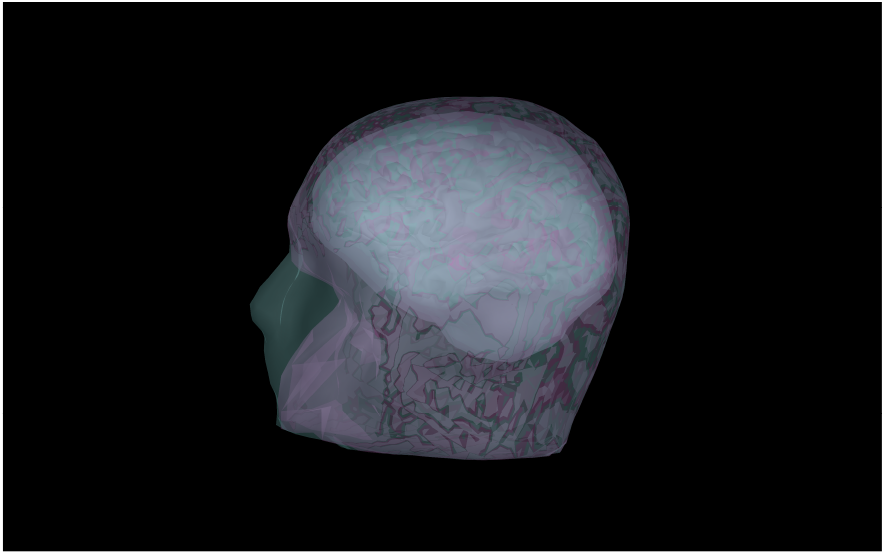
Defaced and non-defaced anatomy of subject 8.

The results indicate that FEM models are significantly more affected by defacing than BEM models. This outcome is expected, as FEM methods incorporate more detailed anatomical information. Although, for this study, only three layers were used when generating the FEM models, the impact of defacing still remains more pronounced compared to BEM models. Future research could investigate how FEM models with additional layers are influenced by defacing.

One limitation of this study is that performing defacing after preprocessing of the full image may often not be in possible practical applications. Publicly available datasets are typically pre-defaced but have not always been processed by a pipeline like FreeSurfer `recon-all`. In this work, defacing was applied post-preprocessing to maintain consistency in the source space between the defaced and non-defaced anatomies and to avoid the need for additional cortical location mappings. However, we expect the resulting underestimation of defacing-induced errors to be minor.

## V. CONCLUSION

The study explores the impact of defacing on EEG source localization, offering a quantitative comparison between head models generated from original and defaced anatomies. The analysis is performed using both the boundary and finite element methods, respectively. The results indicate thatdefacing affects the generated lead field computation and EEG source localization. As such, it is recommended to construct EEG volume conductor models before defacing if accurate source localization is desired.

However, the effects vary across the ten subject cohort, highlighting the need for further investigation. Future studies should employ more advanced FEM methods to obtain more detailed models for a deeper assessment of defacing effects. Additionally, it would be interesting to explore the effect of defacing with real EEG data instead of a simulation. Finally, future studies should investigate the ability of facial reconstruction algorithms to mitigate the effect of defacing. Thus, what is presented in this work sets up a testbed for more in-depth studies and simulations in the future.

## ACKNOWLEDGMENT

The authors thank Nicolas Langer and Tzvetan Popov for initial discussions.

